# Transferability, development of Single Sequence Repeat (SSR) markers and application to the analysis of genetic diversity and population structure of the African fan palm (*Borassus aethiopum* Mart.) in Benin

**DOI:** 10.1101/2020.01.30.926626

**Authors:** Mariano Joly Kpaténon, Valère Kolawolé Salako, Sylvain Santoni, Leila Zekraoui, Muriel Latreille, Christine Tollon-Cordet, Cédric Mariac, Estelle Jaligot, Thierry Beulé, Kifouli Adéoti

## Abstract

In Sub-Saharan Africa, the fan palm *Borassus aethiopum* Mart. is an important non-timber forest product-providing palm that faces multiple anthropogenic threats to its genetic diversity. However, this species is so far under-studied, which prevents its sustainable development as a resource.

The present work is a first attempt at characterizing the genetic diversity of this palm species as well as its spatial structuration in Benin, West Africa. During a first phase we implemented a microsatellite markers-based approach relying on the reported transferability of primers developed in other palm species and found that, in disagreement with previously published results, only 22.5% of the 80 markers tested enabled amplification of African fan palm DNA and polymorphism detection was insufficient. During a second phase, we therefore generated a *B. aethiopum*-specific genomic dataset through high-throughput sequencing and used it for the *de novo* detection of potential microsatellite markers. Among these, 11 enabled polymorphism detection and were further used for analyzing genetic diversity in nine *B. aethiopum* populations.

Our results show that genetic diversity of Beninese fan palm populations is low, with an overall average expected heterozygosity (He) of 0.354. Moreover, the positive values of the fixation index (F) in populations from both the Central (Soudano-Guinean) and the Southern (Guinean) regions suggest limited gene flows. Our analysis show that sampled *B. aethiopum* populations are clustered into two groups, one spanning populations from both the Southern and most of the Central region, and the other including the Central population of Savè (which also has the highest He) and populations from the North.

In light of our results, we discuss the use of inter-species transfer vs. *de novo* development of microsatellite markers in genetic diversity analyses targeting under-studied species. We also suggest future applications for the molecular resources generated through the present study.

## Introduction

Many plant species remain under-studied due to their low economic importance, complicated biology and/or the absence of available genome sequence information. Upon initiating a research project aimed at characterizing the genetic diversity of such a species, researchers may be confronted with the situation that some resources can be found in more or less distantly related taxa. In such cases, the first step is often to assess whether some of these resources, such as molecular markers, can be used to study the new species. Provided that the “source” species display enough genetic similarities to the “target” species and that marker transferability has been previously assessed, this first step may lead to quick progress in a cost-effective manner. In many instances, transferring markers between species is therefore seen as a smarter investment than developing and testing new markers, especially if the initial funding allocated to the project is scarce.

Over the last three decades, molecular markers have been widely used to study genetic variation among and within populations of various plant species [1–10]. Among the different types of markers that are available, microsatellites or Single Sequence Repeats (SSRs) are often selected because they are easy to use and their implementation has low resources (*i.e.* genomic, financial, lab equipment) requirements. As a result, they are markers of choice for the assessment of polymorphism among species, genetic structure within populations, phylogeny reconstruction, genetic mapping, evolutionary analysis, and molecular breeding [11–14]. However, the steps leading to the development of functional SSR markers, namely the initial identification of microsatellite loci, primer selection and assessment of amplification/polymorphism detection, require some prior knowledge of the genome of the target species and may prove to be expensive and time-consuming [13,15]. In order to overcome this difficulty, approaches relying on the transfer of SSR markers between species or genera have therefore been implemented. They have been successful in many instances, as documented across *Prunus* species and among members of the Rosaceae family [16,17]; between species of the *Hevea* genus and to other Euphorbiaceae [18]; among Lamiaceae [19]; among Legumes belonging to the *Vicia* genus [20] and from the *Phaseolus* genus to *Vigna* [21].

The African fan palm *Borassus aethiopum* Mart., also known as ron or toddy palm, is a dioecious species belonging to the Arecaceae family. It is widely distributed across West and Central Africa, where it is present as wild populations. The fan palm is classified as a non-timber forest products (NTFPs)-providing plant [22,23], since the different parts of the plant are used for various purposes by local populations: hypocotyls and fruits for food, fruit odor as shrew repellent, stipe for construction, roots and leaves for traditional medicine, leaves for crafts [24–28]

These multiple uses of products derived from *B. aethiopum* have put a strong anthropogenic pressure on the species, thus contributing to both fragmentations of its populations and their poor natural regeneration [24,29–32]. More specifically, the harvesting of *B. aethiopum* fruits for hypocotyl production and trade has become, over the last two decades, one of the most important household commercial activities associated with this species in Benin, West Africa [33]. Further fragmentation of the species’ habitat has been observed as result of land clearing for agriculture or urban development [32,34,35]. As illustrated through similar examples in the literature [39,40], such phenomena may lead to restricted gene flow and ultimately, to loss of genetic diversity among *B. aethiopum* populations.

There is therefore an urgent need to define a sustainable management policy for *B. aethiopum* populations, in order to ensure its sustainable use. As a consequence, acquiring information on the genetic diversity of the species and on the spatial structuring of its populations is a major touchstone towards defining sustainable management actions. At the time of writing the present article, only a few chloroplastic sequences are publicly available for *B. aethiopum* through NCBI (https://www.ncbi.nlm.nih.gov/search/all/?term=borassus%20aethiopum). By contrast, abundant molecular resources, including genome assemblies or drafts, are available for model palm species such as the African oil palm *Elaeis guineensis* Jacq. [41], the date palm *Phoenix dactylifera* [42–44] and the coconut tree *Cocos nucifera* [42,43]. In each of these three palm species, large numbers of SSR markers have been identified and for a fraction of them, cross-species and cross-genera transferability tests among species belonging to the Palmaceae family have been performed [47–53]. In several instances [48–51,53], these tests included samples from the Asian relative of *B. aethiopum, B. flabellifer*.

In the present study, we describe how we first attempted to use SSR markers that had been identified in these other palm species for the analysis of genetic diversity in *B. aethiopum*. Then, in a second phase, we show how we performed a low-coverage sequencing of the fan palm genome in the aim of developing the first set of specific SSR markers targeting this species. We then used these to assess the genetic diversity of *B. aethiopum* populations in Benin, as a preliminary step towards more comprehensive studies.

## Material and Methods

### Plant material sampling and DNA extraction

Nine distinct populations of *B. aethiopum* separated by at least 50 km were selected from the three main climatic regions that are encountered in Benin (Fig 1): the Sudanian region in the North (four populations), the Sudano-Guinean region in the Centre (three populations) and the Guineo-Congolian region in the South (two populations). Additionally, among sampled populations, three were located in protected areas and six in farmlands. Within each population, we sampled young leaves from 20 male and female adult trees that were separated by at least 100 m, and stored them in plastic bags containing silica gel until further processing. The complete list of leaf samples and their characteristics is available in S1 Table.

**Fig 1.**
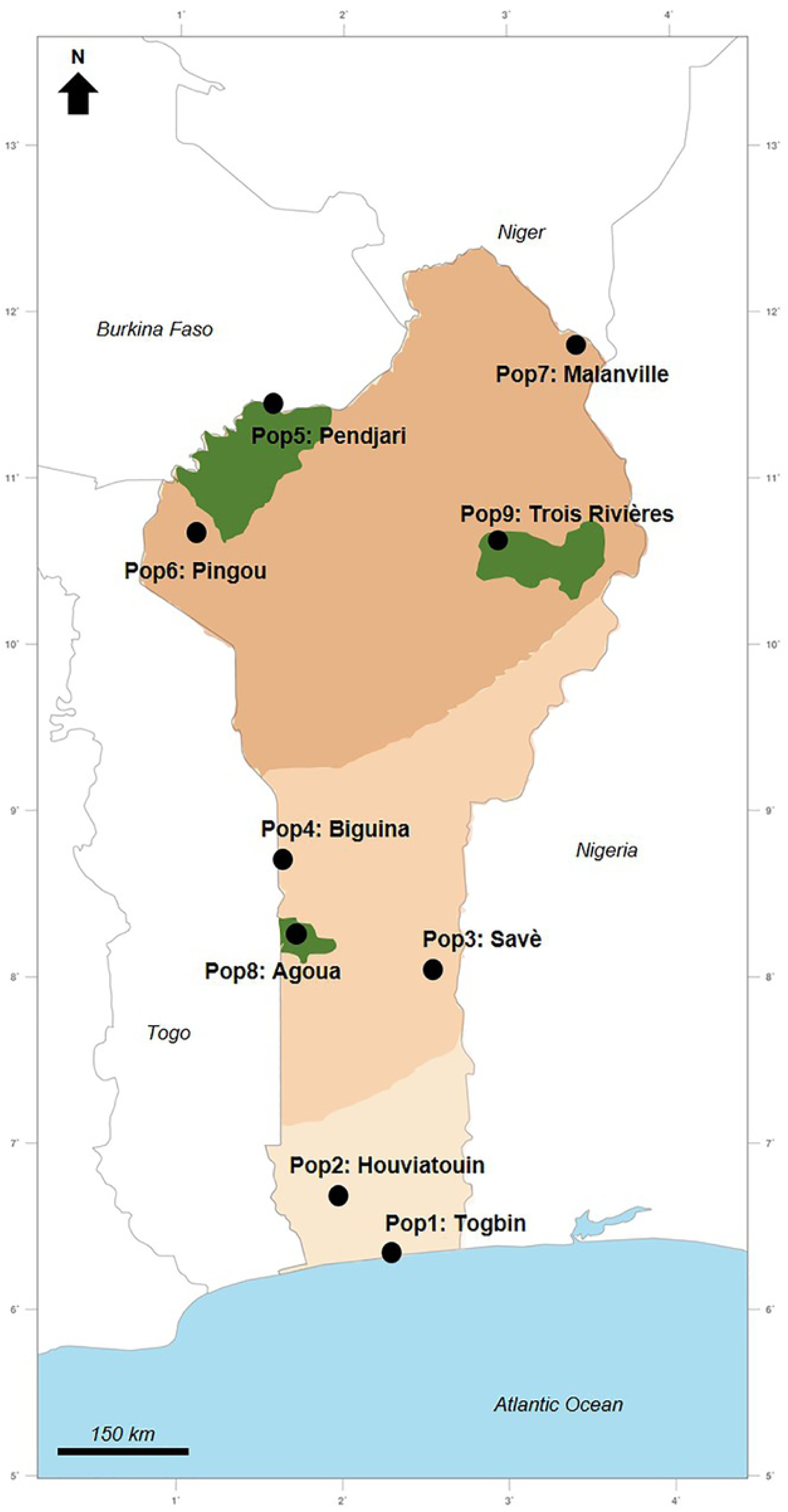
Geo-climatic distribution of the Beninese *Borassus aethiopum* populations used in this study. Collection sites for each of the nine populations sampled are indicated; information for individual samples are available in S1 Table. The three main geo-climatic regions of Benin are (from the lighter- to the darker-colored): Guineo-Congolian, Sudano-Guinean and Sudanian, respectively. Adapted from a map by the GingkoMaps project (http://www.ginkgomaps.com/), distributed under a Creative Commons Attribution (CC-BY) 3.0 license (https://creativecommons.org/licenses/by/3.0/).

Genomic DNA was extracted from 250 mg of leaves ground to powder under liquid nitrogen using the Chemagic DNA Plant Kit (Perkin Elmer, Germany), according to the manufacturer’s instructions on a KingFisher Flex™ (Thermo Fisher Scientific, USA) automated DNA purification workstation. Final DNA concentration was assessed fluorometrically with the GENios Plus reader (TECAN) using bis-benzimide H 33258 (Sigma-Aldrich) as a fluorochrome.

### Transferability of palms microsatellite markers: selection and amplification

We selected a total of 80 SSR markers from previous studies: 44 were developed on *Phoenix dactylifera*; 25 were identified in *Elaeis guineensis* and showed successful amplification on *Borassus flabellifer* and *P. dactylifera* DNA; and 11 from *Cocos nucifera*. The respective sequences and origins of these different primer sets are displayed in Table 1.

**Table 1:**
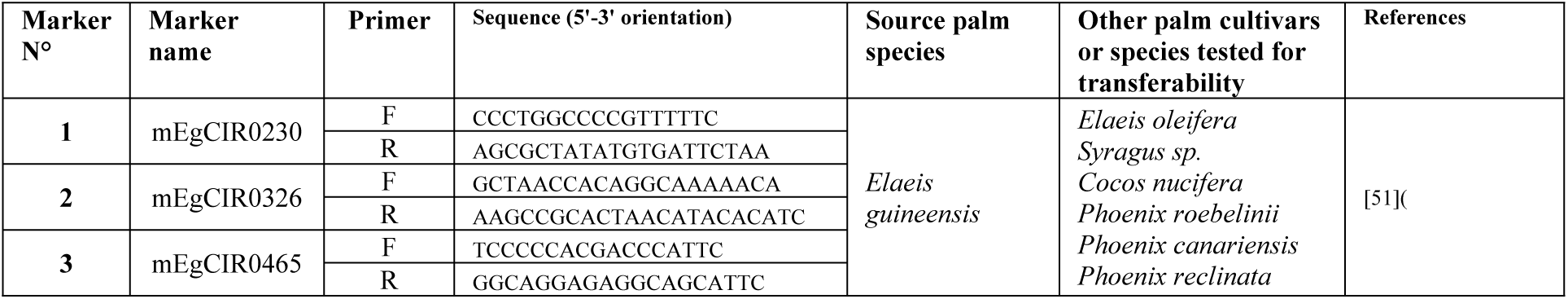
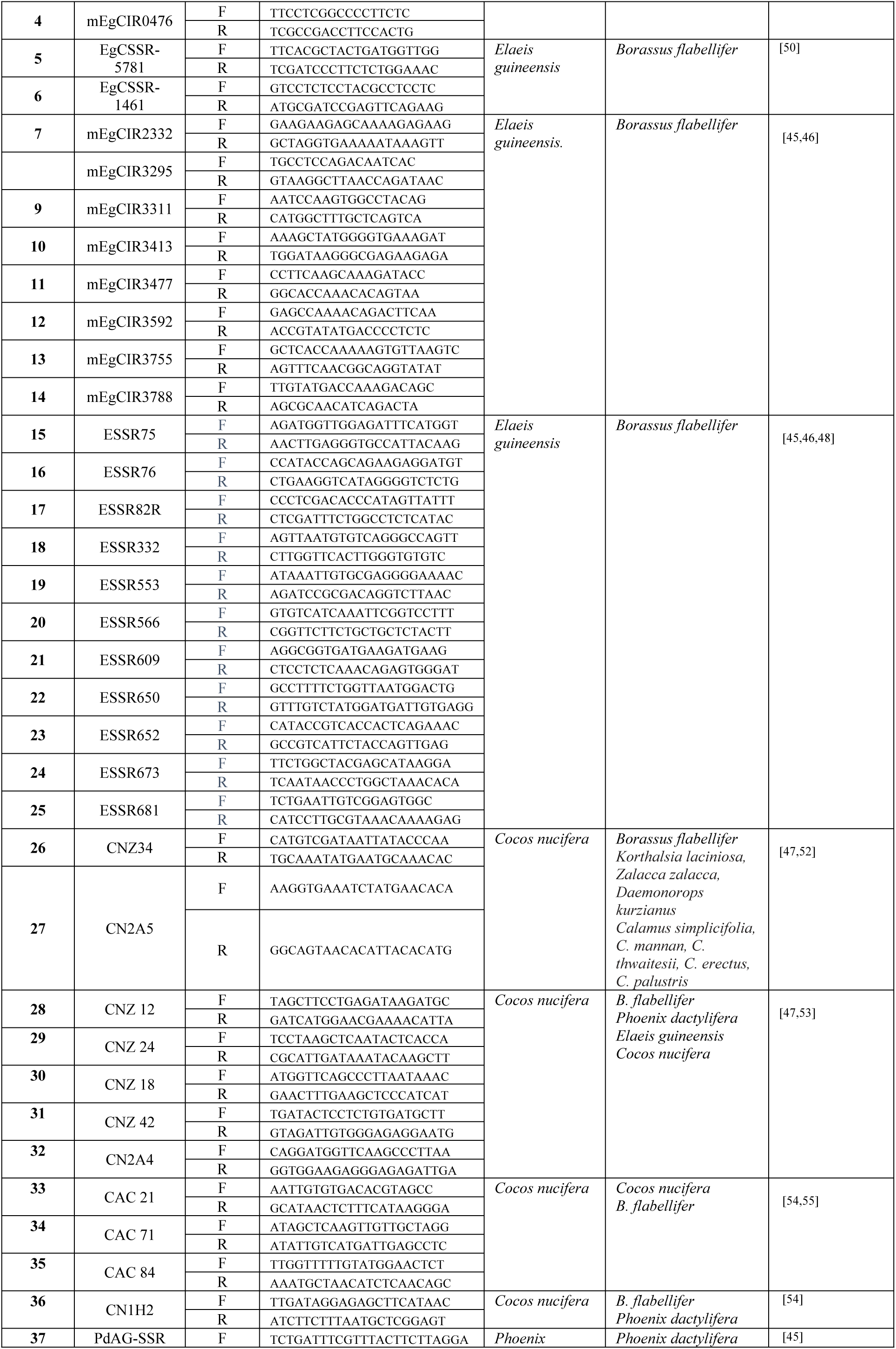
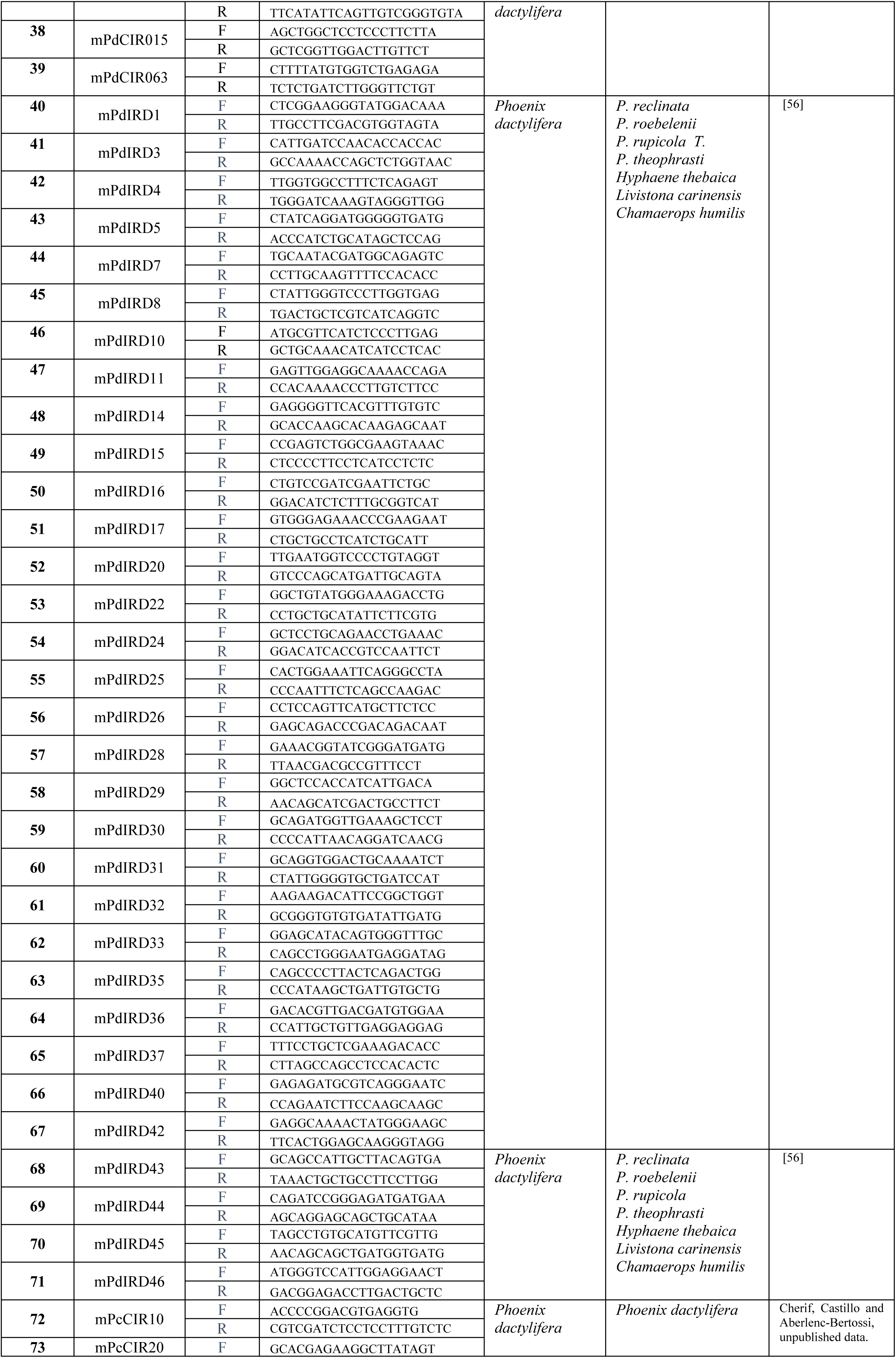
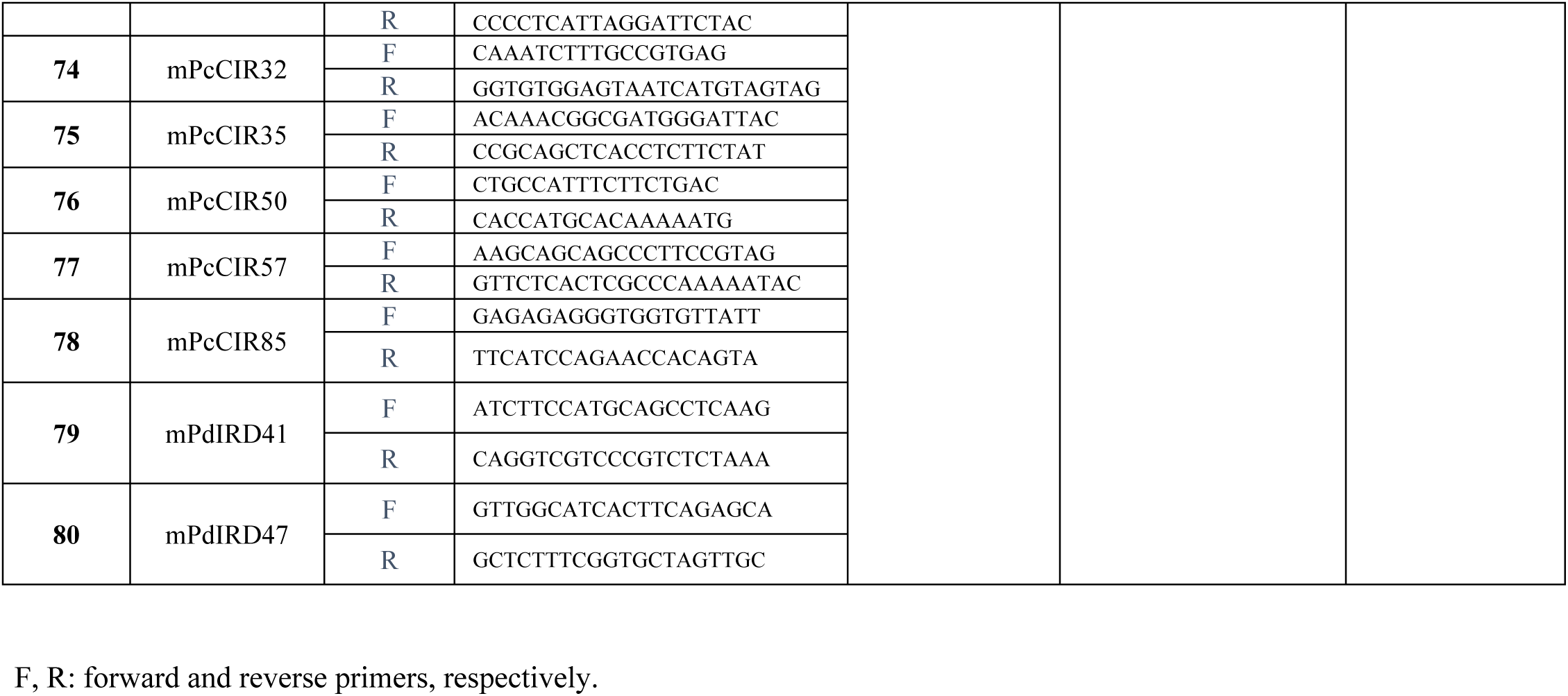
Characteristics of the palm SSR markers tested for transferability on *B. aethiopum*.

Transferability of the 80 palm SSR markers was assessed on a representative subset of 20 of *B. aethiopum* individuals sampled in the different populations, plus 4 positive controls from each source species for these markers (*P. dactylifera, C. nucifera, E. guineensis*). Microsatellite amplification was performed with a modification of the M13-tailed Primers protocol (Boutin-Ganache et al., 2001) adapted to the use of fluorescent labelling. The PCR reaction was performed on 20 ng of leaf DNA and a mix of 1X PCR buffer, 200 μM dNTP, 2 mM MgCl2, 0.4 pmol M13-tailed forward primer fluorescently labeled in 5’ with FAM, HEX or TAMR, 4 pmol reverse primer, and 0.5 U of *Taq* polymerase (Sigma), in a final volume of 20 µl. The following program was used: 3 min of initial denaturation at 95°C, followed by 35 cycles of 30 s at 95°C, 30 s at 50°C and 72°C for 1 min and a final extension at 72°C for 5 min. The resulting amplification products were then diluted to 1/10^th^ mixed with 0.5 µL of an internal size standard (GeneScan 500 ROX, Thermo Fisher Scientific) and denatured for 5 minutes at 94°C prior to capillary electrophoresis (Applied Biosystems 3500 Genetic Analyzer, Thermo Fisher Scientific).

### *De novo* identification of microsatellite loci in the *B. aethiopum* genome, marker selection and diversity analysis

One *B. aethiopum* leaf sample was randomly selected and genomic DNA purification was performed according to the protocol of [57]. This DNA extract was then used for the construction of an Illumina paired-end library, as described in [58], before high-throughput sequencing on a MiSeq v3 Illumina platform. Demultiplexing of the raw data output was performed using the Maillol script (https://github.com/maillol/demultadapt), with a 0-mistmatch threshold. Adapters were eliminated using Cutadapt v1.10 (Martin, 2011) (http://code.google.com/p/cutadapt/) with the following parameters: overlap length = 7, minimum length = 35 and quality = 20. High-quality reads (Q > 30) were filtered using the following script: https://github.com/SouthGreenPlatform/arcad-hts/blob/master/scripts/arcad_hts_2_Filter_Fastq_On_Mean_Quality.pl and the resulting filtered reads were deposited into GenBank SRA under accession number PRJNA576413. Paired-end reads were then merged using FLASH v1.2.11 (https://github.com/SouthGreenPlatform/arcad-hts/blob/master/scripts/arcad_hts_3_synchronized_paired_fastq.pl). Finally, microsatellite motif detection and specific primer design were carried out after elimination of redundant sequences using the QDD v3.1.2 software [59] with default settings.

Using selected primer pairs, test amplifications were performed with two randomly selected fan palm DNA samples, then primers showing successful amplification were further tested for polymorphism detection among seven randomly selected DNA samples. The M13 Tailed Primers protocol described previously was used, with the following program: 3 min of initial denaturation at 95°C, followed by 35 cycles of 30 s at 95°C, 30 s at 55°C and 72°C for 1 min and a final extension at 72°C for 5 min. PCR products visualization was performed as previously indicated. Finally, the primer pairs enabling successful amplification of polymorphic, mono-locus bands were used for the analysis of genetic diversity among the complete set of 180 *B. aethiopum* individuals with the same conditions.

### Data analysis

Amplification products were scored using the GeneMapper software V3.7 and only unambiguous amplification products were considered for data analysis. Genetic parameters such as total number of alleles, allelic frequency, expected heterozygosity (He), observed heterozygosity (Ho), were calculated for each locus and each population in the GenAIEx software Version 6.502 [60]. The F-statistics analysis assessing genetic differentiation and the Analysis of MOlecular VAriance (AMOVA) for estimation of genetic differentiation within and among populations were performed with the same software.

A Principal Coordinates Analysis (PCoA) was also performed using GenAIEx software to enable the visualization of genetic variation distribution across the individuals under study. We used the STRUCTURE software version 2.3.4 [62] for the determination of the most probable number of clusters for population structure (K value). Using the admixture model, eight simulations were performed for each inferred K value, with a running length composed of 300,000 burning periods and 50,000 Markov chain Monte Carlo (MCMC) iterations to allocate accessions to different populations. The output from this analysis was then used as input in the Structure HARVESTER online program to determine the exact [62]. Based on this value, a clustering analysis of the studied populations was performed and using the genetic distance matrix obtained from previous analysis, a dendrogram was constructed with the DendroUPGMA program accessible online at http://genomes.urv.cat/UPGMA/ [63]

## Results

### Assessment of palm SSR markers transferability to *B. aethiopum* and evaluation of their capacity for characterizing genetic diversity

Of the 80 microsatellite markers that have been selected from three palm species and tested for amplification on African fan palm DNA, 18 (22.5 %) generate amplification products (Table 2). No amplification is observed using the 11 *C. nucifera* markers, whereas 7 (15.9 %) and 11 (44%) of the *P. dactylifera* and *E. guineensis* markers, respectively, show a successful amplification. None of the amplification products generated with date palm primers display genetic polymorphism in our *B. aethiopum* test panel. Among oil palm-derived SSR markers however, two, namely ESSR566 and ESSR652, display polymorphism. However, it must be noted that the ESSR566 primer pair amplifies two distinct loci, ESSR566A and ESSR566B.

**Table 2:** Summary of SSR markers transferability assessment.

| Species of origin | Number of SSR markers | Number of successful amplification (% of markers) | Number of polymorphic amplicons (% of amplifications) |
| --- | --- | --- | --- |
| *Cocos nucifera* | 11 | 0 (0) | 0 (0) |
| *Phoenix dactylifera* | 44 | 7 (15.9) | 0 (0) |
| *Elaeis guineensis* | 25 | 11 (44.0) | 2 (18.2) |
| **Total** | **80** | **18 (22.5)** | **2 (11.1)** |

Overall, during this phase of the study we detect polymorphism in our *B. aethiopum* test panel with only 2 (11.1% of successfully amplified markers, 2.5% of total) of the palm SSR primer pairs that have been assayed. Only one of these markers proves to be both polymorphic and monolocus in the African fan palm, and might therefore be used for studying genetic diversity in this species.

### *De novo* identification of microsatellite sequences in the *B. aethiopum* genome and assessment of potential SSR markers

In order to enable a more precise evaluation of genetic diversity among *B. aethiopum* populations, we developed specific *B. aethiopum* markers from *de novo* sequencing data. A total of 23,281,354 raw reads (average length 250 bp) have been generated from one MiSeq run. Raw sequence reads have been trimmed and generated 21,636,172 cleaned-up reads, yielding 493,636 high-quality reads after filtering (Q > 30) from which 216,475 contigs have been assembled.

From this latter output, the QDD software identifies a total of 1,630 microsatellite loci (see S2 Table), of which 81.41 % are perfect (*i.e.* repeat size 4 bp or smaller and repeat number 10-20). Among these, 83.86 % of loci are composed of di-nucleotidic repeat units, 13.06 % of tri-nucleotidic units, 2.39 % of tetra-nucleotidic repeats and 0.67 % of repeats with five nucleotides and over. From these, we have selected SSR markers composed of di- (AG) or tri-nucleotide repeats, using the following criteria for specific amplification of easily scorable bands: primer lengths ranging from 18 to 22 bp, annealing temperatures 55–60°C and predicted amplicon sizes 90-200 bp.

The characteristics of the 57 selected primer pairs and the results of the test amplifications are presented in Table 3. Successful amplification of *B. aethiopum* DNA is obtained in most cases (94.7%). However, 34 of the putative markers tested (63.0% of amplifying ones) show no polymorphism. The remaining 20 putative markers are polymorphic and among them, nine correspond to multiple loci. As a result, 11 putative African fan palm SSR markers (representing 20.4% of successful amplifications and 55.0% of polymorphic markers in our study) are both polymorphic and mono-locus in our amplification test panel and may therefore be used for further analyses.

**Table 3:**
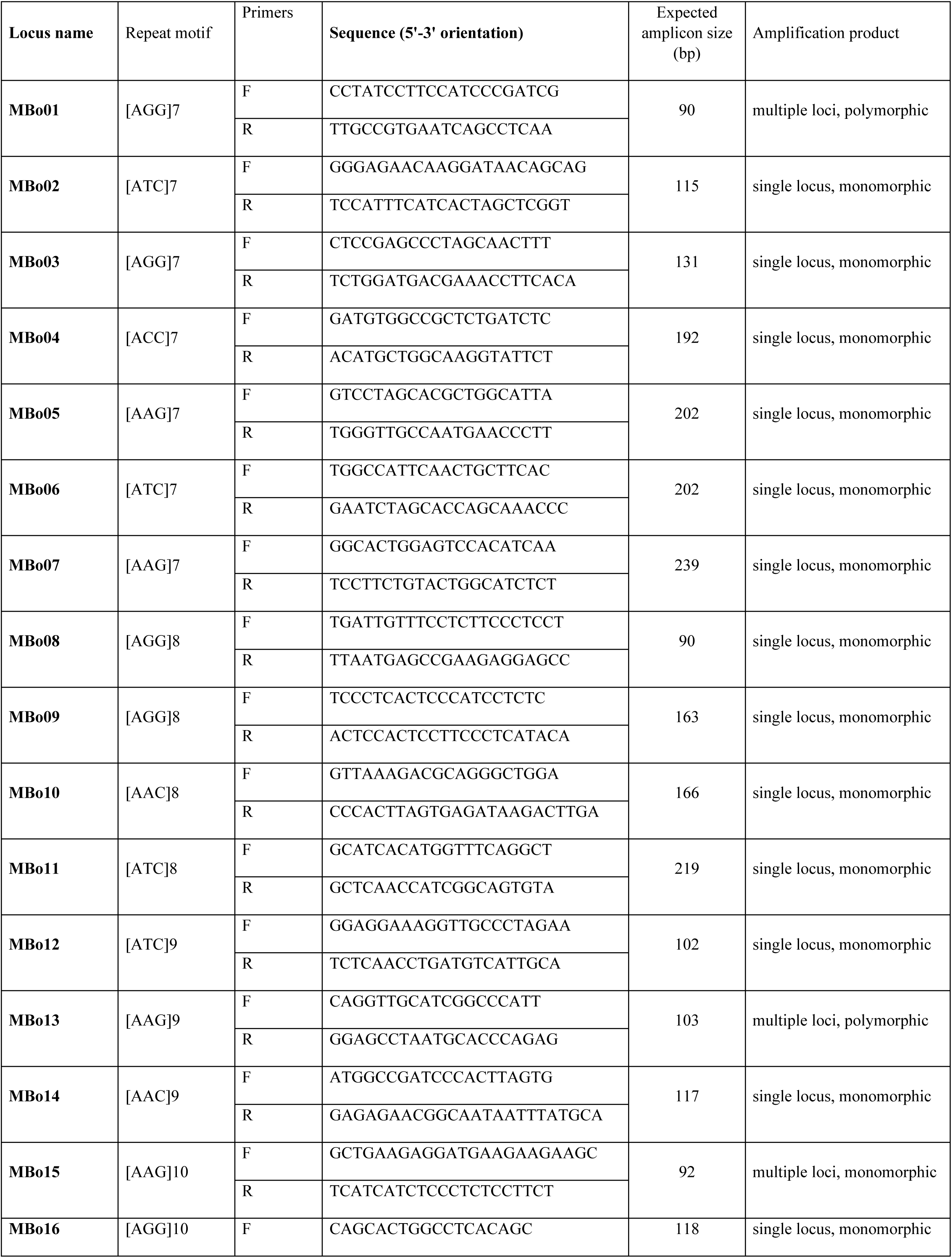
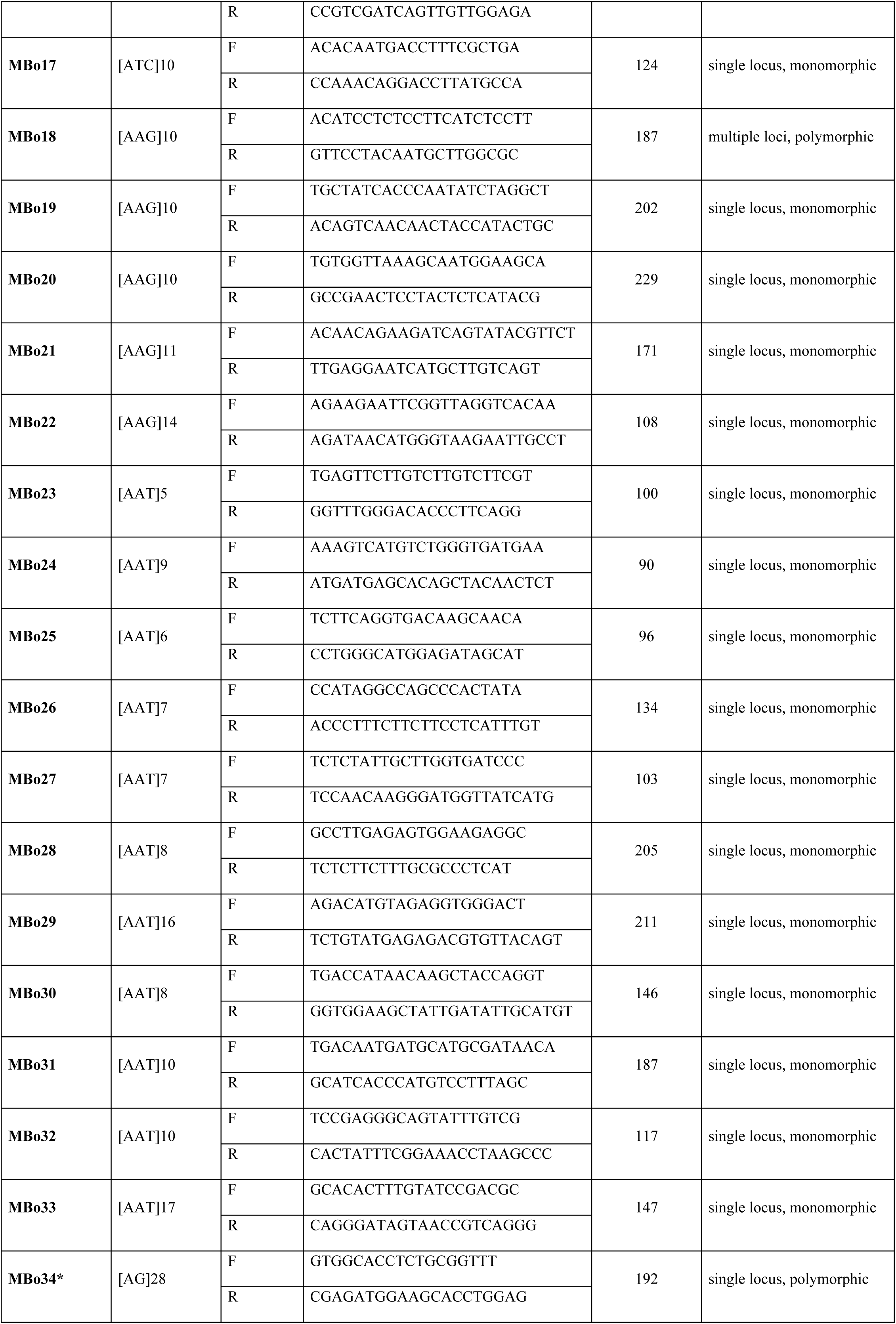
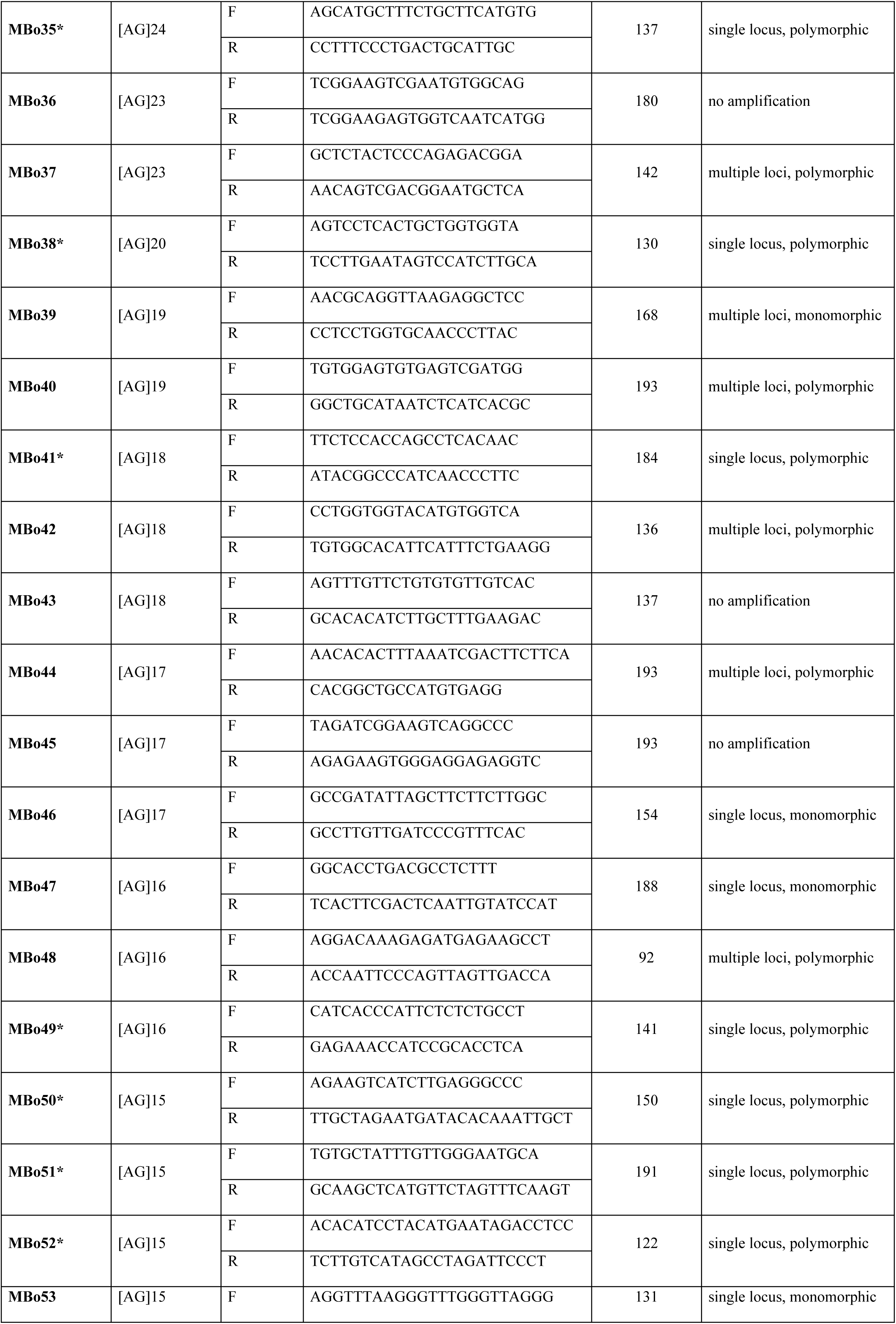
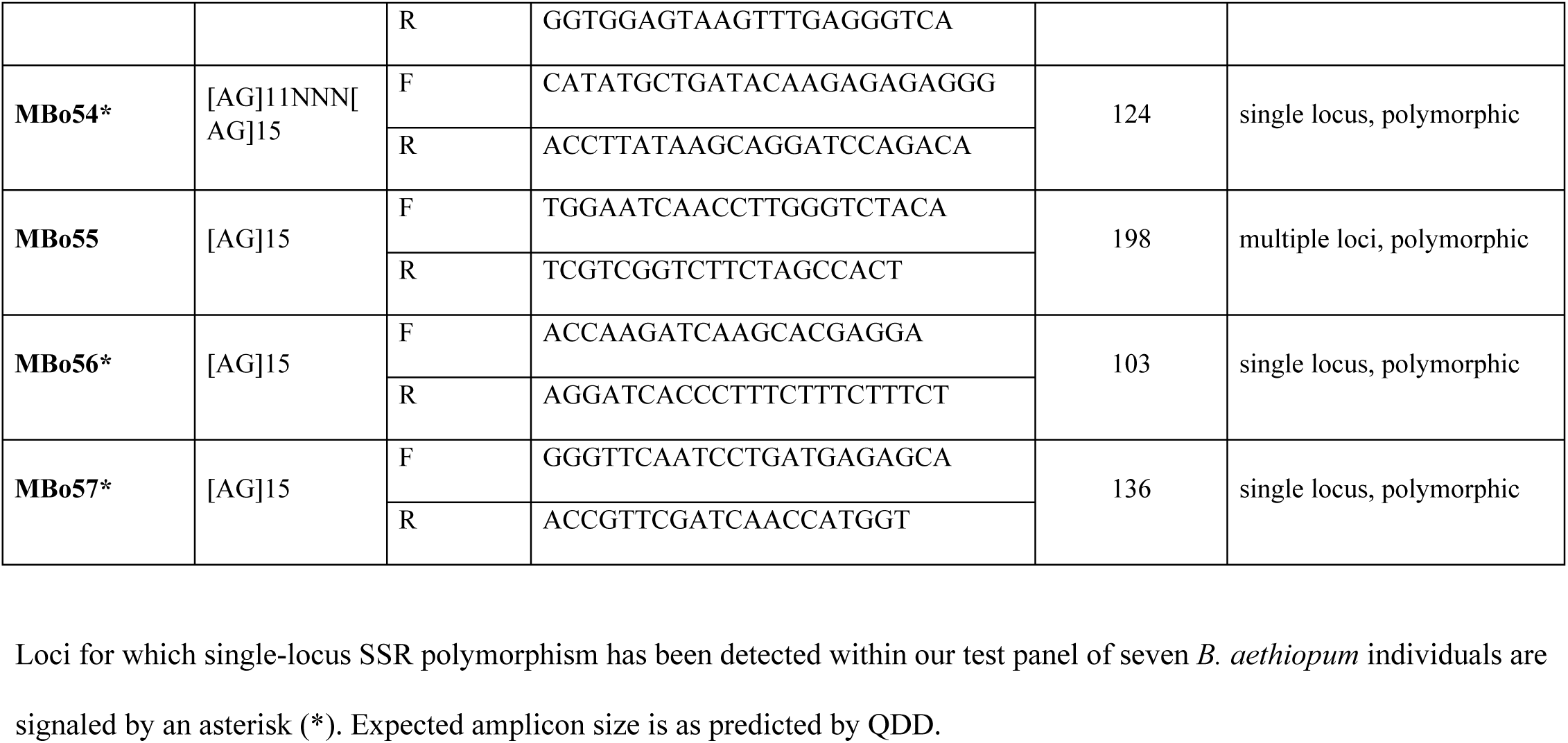
List of selected primer pairs targeting putative *B. aethiopum* microsatellite loci and assessment of their polymorphism detection ability

### Microsatellite-based characterization of the genetic variation within *B. aethiopum* populations of Benin

The set of 11 *B. aethiopum*-specific SSR markers identified in the previous step has been used for the characterization of genetic diversity in our full panel of nine populations (180 individuals) distributed across Benin. Among our sample set, the number of alleles per microsatellite locus ranges from 2 for locus Mbo41 to 6 for loci Mbo34, Mbo35 and Mbo50, with an average value of 4.27, whereas expected heterozygosity (He) values range from 0.031 (locus Mbo56) to 0.571 (locus Mbo35; Table 4). Using these markers, the analysis of genetic diversity (Table 5) shows that the percentage of polymorphism detected at the microsatellite loci investigated ranges from 72.73% (populations of Togbin and Malanville) to 90.91% (populations of Savè, Agoua, Pendjari, Pingou and Trois Rivières), with a mean value of 84.85%. With the exception of the Savè, Hounviatouin and Malanville populations, 1 to 3 private alleles of the targeted microsatellite loci are observed in most populations. Regarding the genetic parameters, the number of effective alleles (Ne) ranges from 1.447 to 2.069 with an average number of 1.761. He values range from 0.263 (Hounviatouin) to 0.451 (Savè) with an average value of 0.354 whereas the observed heterozygosity (Ho) varied from 0.234 (Togbin) to 0.405 (Pingou) with an average value of 0.335. Negative values of Fixation index (F) are obtained for the populations of Pingou, Malanville and Trois rivières whereas positives F values are observed in all other populations investigated, indicating a deficit of heterozygosity in the latter.

**Table 4:**
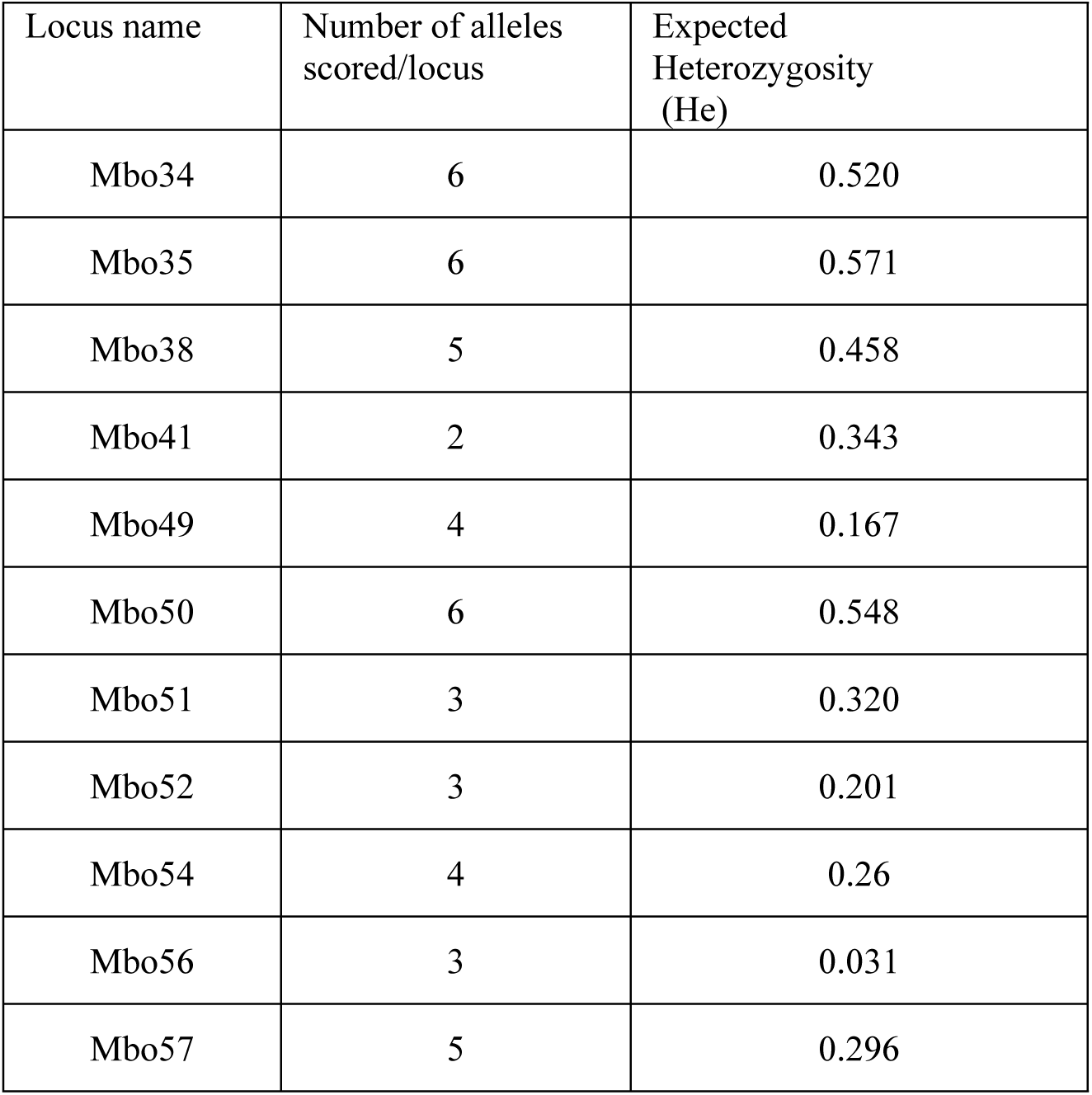
Characteristics of 11 polymorphic microsatellites markers used for genetic diversity analysis of *B. aethiopum*

**Table 5:**
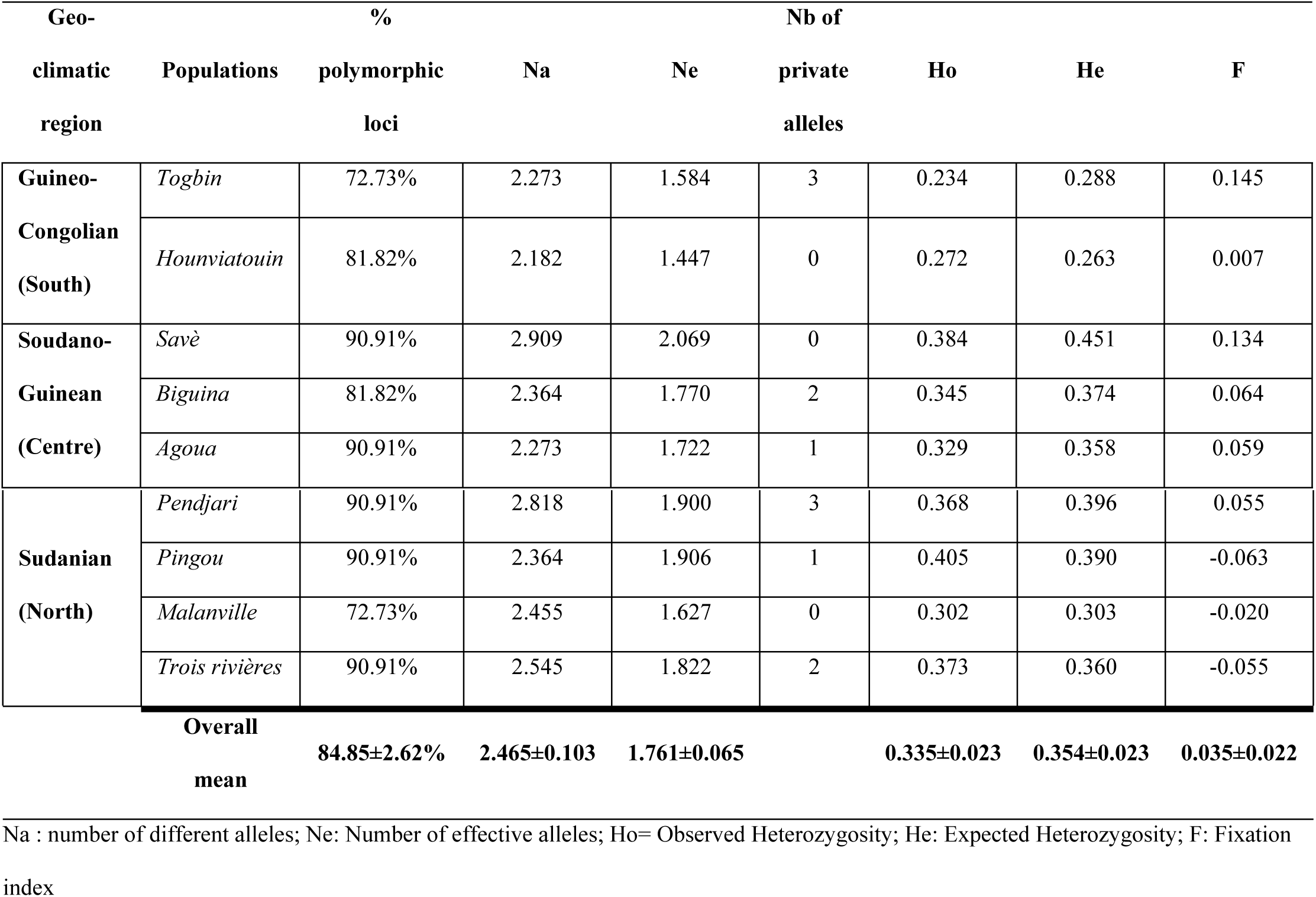
Mean diversity parameters for each of the nine *B. aethiopum* populations.

### Genetic structure of the *B. aethiopum* populations under study

The calculation of Nei’s genetic distance among populations (Table 6) shows values ranging from 0.073, as observed between Togbin and Hounviatouin (Guineo-Congolian region), to 0.577 between Togbin (Guineo-Congolian region) and Trois Rivières (Sudanian region). Overall, genetic distances between the fan palm populations under study are lowest within the same region, with the lowest genetic distances among populations of Savè, Pendjari, Pingou, and Trois Rivières which are all located in the Northern part of the country. One interesting exception is the Centre (Guineo-Sudanian) region of Benin, where we find that the most genetically distant population from Savè is the one collected within the Agoua forest reserve (0.339). Surprisingly, Savè displays its highest genetic identity value when compared to the other two populations sampled in protected areas, namely Pendjari (0.870) and Trois Rivières (0.882) which are both located in the Sudanian region. This is an unexpected finding considering the important geographic distances that are involved.

**Table 6:** Pairwise Population Matrix of Nei’s genetic distance and genetic identity values.

|  | **Togbin** | **Hounviatouin** | **Savè** | **Biguina** | **Agoua** | **Pendjari** | **Pingou** | **Malanville** | **Trois Rivières** |
| --- | --- | --- | --- | --- | --- | --- | --- | --- | --- |
| **Togbin** | - | 0.073 | 0.477 | 0.253 | 0.337 | 0.517 | 0.494 | 0.487 | 0.577 |
| **Hounviatouin** | 0.929 | - | 0.419 | 0.110 | 0.215 | 0.435 | 0.317 | 0.375 | 0.535 |
| **Savè** | 0.621 | 0.658 | - | 0.270 | 0.339 | 0.140 | 0.265 | 0.238 | 0.126 |
| **Biguina** | 0.776 | 0.896 | 0.763 | - | 0.152 | 0.241 | 0.161 | 0.186 | 0.316 |
| **Agoua** | 0.714 | 0.806 | 0.713 | 0.859 | - | 0.408 | 0.304 | 0.359 | 0.490 |
| **Pendjari** | 0.596 | 0.647 | 0.870 | 0.786 | 0.665 | - | 0.167 | 0.108 | 0.103 |
| **Pingou** | 0.610 | 0.728 | 0.767 | 0.851 | 0.738 | 0.846 | - | 0.174 | 0.175 |
| **Malanville** | 0.614 | 0.688 | 0.788 | 0.831 | 0.699 | 0.898 | 0.841 | - | 0.145 |
| **Trois Rivières** | 0.561 | 0.585 | 0.882 | 0.729 | 0.613 | 0.902 | 0.840 | 0.865 | - |

Above the diagonal: Nei's genetic distance; below: genetic identity.

A similar structuration of genetic distances emerges from the analysis of pairwise population genetic differentiation (Fst) (Table 7), suggesting genetic differentiation according to geographic distances between populations, with the notable exception of the lower genetic differentiation between palms from Savè and those from either one of the forest reserves in the Northern region.

**Table 7:** Pairwise populations Fst value.

|  | **Togbin** | **Hounviatouin** | **Savè** | **Biguina** | **Agoua** | **Pendjari** | **Pingou** | **Malanville** | **Trois Rivières** |
| --- | --- | --- | --- | --- | --- | --- | --- | --- | --- |
| **Togbin** | 0.000 |  |  |  |  |  |  |  |  |
| **Hounviatouin** | 0.072 | 0.000 |  |  |  |  |  |  |  |
| **Savè** | 0.233 | 0.221 | 0.000 |  |  |  |  |  |  |
| **Biguina** | 0.168 | 0.086 | 0.145 | 0.000 |  |  |  |  |  |
| **Agoua** | 0.215 | 0.153 | 0.157 | 0.105 | 0.000 |  |  |  |  |
| **Pendjari** | 0.247 | 0.212 | 0.077 | 0.120 | 0.188 | 0.000 |  |  |  |
| **Pingou** | 0.252 | 0.181 | 0.138 | 0.103 | 0.169 | 0.100 | 0.000 |  |  |
| **Malanville** | 0.301 | 0.246 | 0.149 | 0.121 | 0.197 | 0.072 | 0.119 | 0.000 |  |
| **Trois Rivières** | 0.285 | 0.279 | 0.076 | 0.178 | 0.224 | 0.073 | 0.104 | 0.107 | 0.000 |

Our analysis of molecular variance (AMOVA; Table 8) shows that within-population variation underlies the major part (53%) of total variance, whereas among-populations and among-regions variations explain variance to a similar extent (23 and 24%, respectively).

**Table 8:** AMOVA results.

| **Source** | **df** | **SS** | **MS** | **Est. Var.** | **% total variance** | **P value** |
| --- | --- | --- | --- | --- | --- | --- |
| Among Regions | 2 | 309.407 | 154.704 | 1.944 | 24% | <0.001 |
| Among Populations | 6 | 254.302 | 42.384 | 1.903 | 23% | <0.001 |
| Within Populations | 171 | 739.100 | 4.322 | 4.322 | 53% | <0.001 |
| Total | 179 | 1302.809 |  | 8.169 | 100% |  |

df=degree of freedom, SS=sum of squares, MS mean squares, Est. var.=estimated variance

In accordance with results from both the analysis of genetic distances and the AMOVA, the Principal Coordinates Analysis (PCoA) of our 180 individual *B. aethiopum* samples shows that the first axis (accounting for 24% of total variation) distinguishes roughly between two main groups of populations (Fig 2). Likewise, the Bayesian analysis of our data indicates an optimal value of K=2 for the clustering of the studied populations into two groups (Fig 3): one group that includes palms belonging to the populations of Togbin and Hounviatouin from the Southern part of the country, as well as most of the palms from Biguina and Agoua from the Western (Togolese) border of the Centre region; and one group composed of the majority of the palms collected in Savè (Eastern part of the Centre region) and palms from the Northern populations of Pendjari, Pingou, Malanville and Trois Rivières. The dendrogram derived from the UPGMA analysis of our data further shows that, within these two main groups, subgroups can be defined based on geo-climatic regions, Savè being the only exception to this general trend (Fig 4).

**Fig 2.**
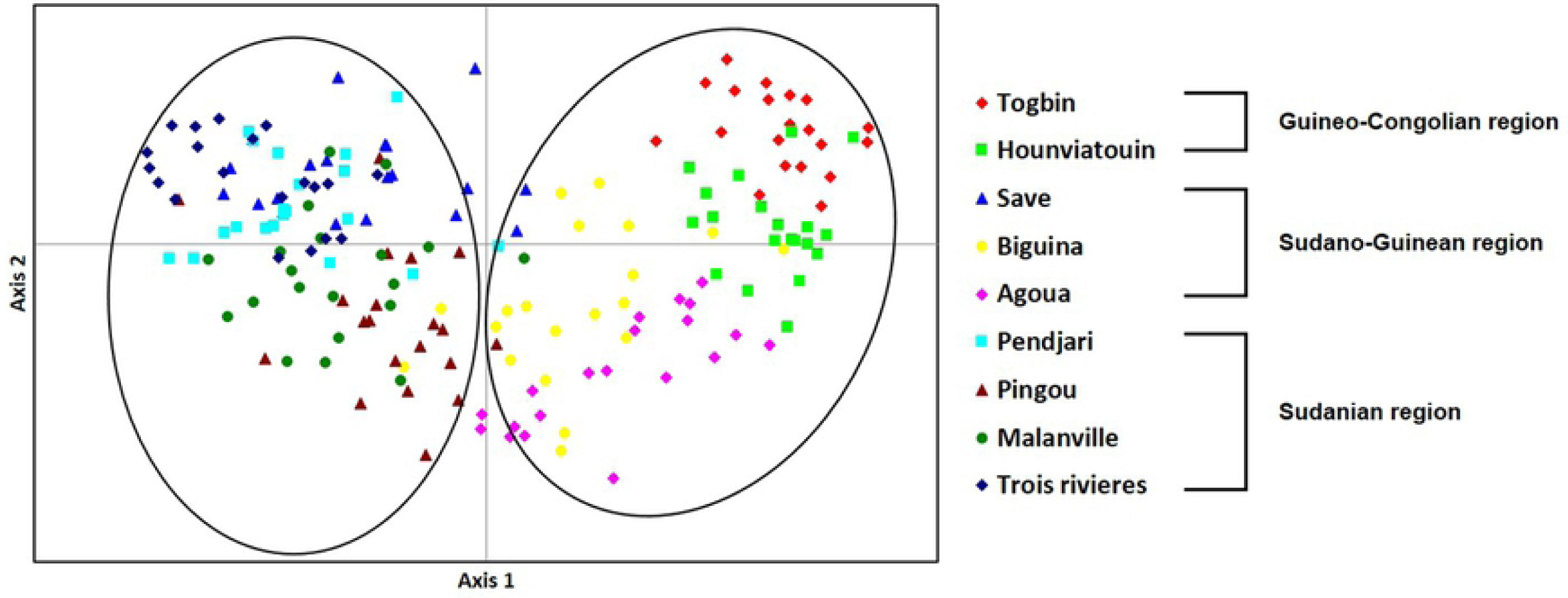
Principal Coordinates Analysis (PCoA) of individual *Borassus aethiopum* samples.

**Fig 3.**
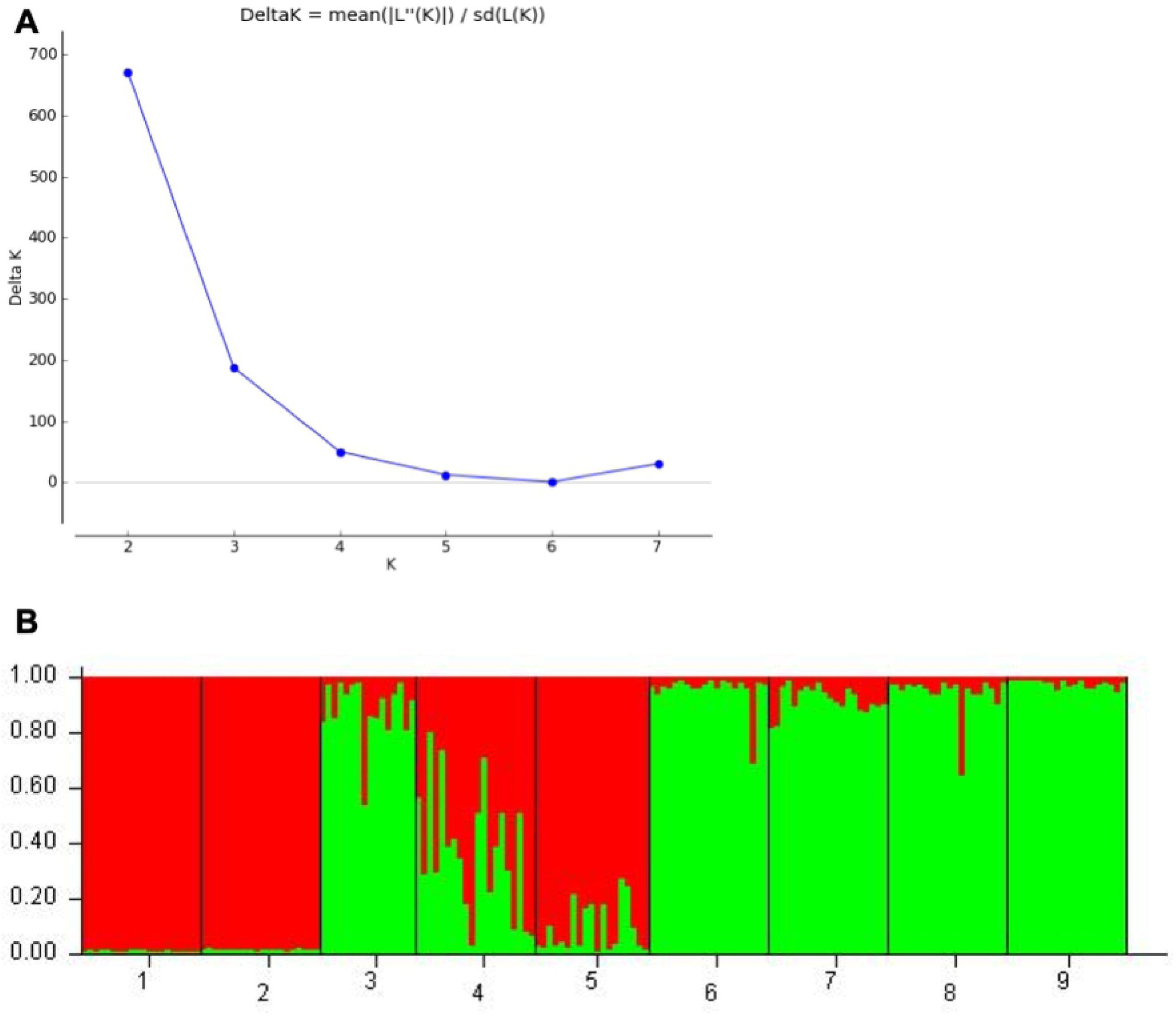
Bayesian cluster analysis. A: Determination of the optimal value of K from Structure Harvester. B: Bayesian STRUCTURE bar plot analysis of Beninese *B. aethiopum* samples with K=2. Red: group 1; green: group 2. Populations are numbered as in S1 Table and displayed along the horizontal axis.

**Fig 4.**
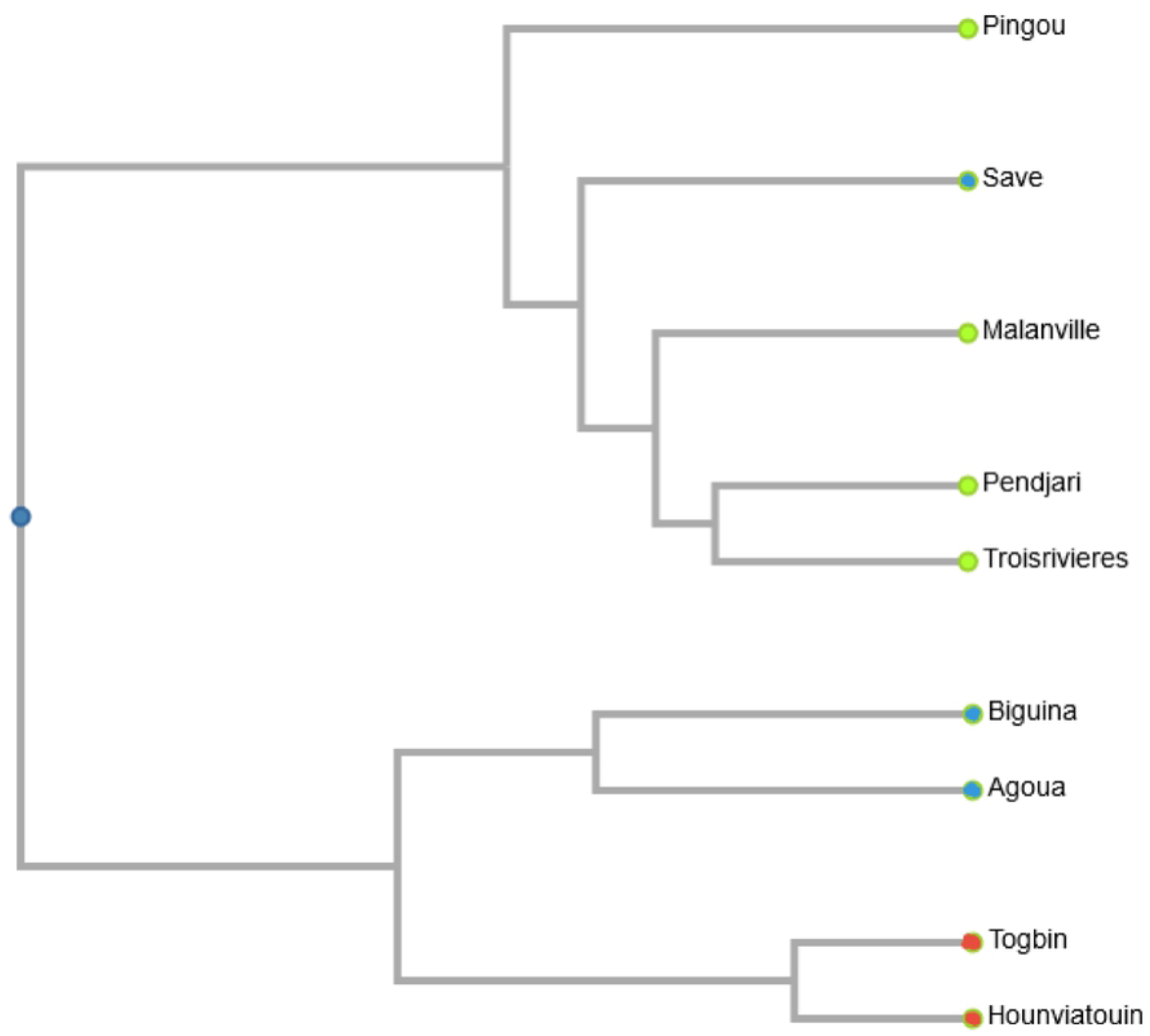
Genetic relationships among Beninese *B. aethiopum* populations. Branch ends with identical colors represent populations from the same geo-climatic region.

## Discussion

In flowering plant, the efficiency of cross-species transfer of SSR markers is highly variable among taxa, especially when important differences in genome complexity exist between the marker source and the target [64]. Nevertheless, this method has been used successfully for accelerating the analysis of genetic diversity in many plant species, including palms [13,68–70] In the present study, we find that the transferability rate of microsatellite markers developed in other palms genera to the African fan palm, *i.e.* their ability to successfully amplify genomic DNA from the latter species, is very low. Indeed, among the 80 primer pairs designed on either oil palm, date palm or coconut palm, we observe that only 22.5% produce amplicons from *B. aethiopum*. This percentage is very low when compared to both the inter-species and inter-genera transferability rates that have been found in similar studies targeting other palm species: from 17 to 93% in a panel of 32 palm species [50], 75% from *E. oleifera* to *E. guineensis* [67], 86% between the wooly jelly palm *Butia eriospatha* and related species *B. catarinensis* [68] and up to 100% in the licuri palm *Syagrus coronata* [70]. When considering other plant families, our transferability rate is also markedly lower than both the average rate of 50% found by [70] within the *Glycine* genus and among Legumes genera, and the overall rate of 35.2% calculated by [71] for within-family transferability among Gymnosperms and Angiosperms. The low transferability rate in our study might be explained in part by the fact that we used markers originating from genomic sequences. Indeed, as pointed out by [72], such markers have a lower transferability rate when compared to Expressed Sequence Tags (ESTs)-derived microsatellites due to the higher inter-species sequence variability within non-coding *vs.* coding sequences. Similarly, it is plausible that differences in genome size and complexity among palm species and genera account for our difficulty to identify palm SSR markers that successfully amplify in *B. aethiopum*. As a matter of fact, the size of the *B. aethiopum* genome, as determined by flow cytometry (1C = 7.73 Gb; Jaume Pellicer, unpublished data), is 3.2 to 11.5 times larger than those of the microsatellite source species used in the present study: the date palm genome is estimated to be 671 Mb [40] whereas the oil palm genome is 1.8-1.9 Gb [41,74] and the coconut genome is 2.42 Gb [46]. Most likely, these differences in genome sizes among related diploid plant species rely on differences in Transposable Element contents and associated structural variations such as copy number variants and homologous recombinations [75], which might eventually affect the cross-species amplification ability of SSR primers. The illustration of such a mechanism working at the intra-genus level has been provided by cultivated rice species *Oryza sativa* and its wild relative *O. australiensis* [76]. More generally, gaining a better understanding of genome structures within the *Borassus* genus could also help reconcile our results with previous published reports of successful transfer of SSR markers developed from other palm sources to *Borassus flabellifer* (see references cited in Table 1). Indeed, since the genome size of *B. flabellifer* is only marginally smaller than that of *B. aethiopum* (7.58 Gb; Jaume Pellicer, unpublished data), significant differences in genome composition may be underlying the lack of SSR transferability between both species.

In any case, from the low number of successfully transferred microsatellite markers we could only identify one displaying polymorphism in our fan palm test panel, making it impossible to rely on for analysis of genetic diversity. Still, the fact that so little microsatellite polymorphism (2 out of 18 amplifying primer pairs: 11.1%) could be detected in this subset of 20 palms sampled across different locations throughout Benin is somewhat surprising and its reasons remain to be elucidated. In addition to possibly being a symptom of habitat fragmentation and low gene flow between populations, this low diversity might also result from the extremely long juvenile phase that has been attributed to this palm species, for which authors have reported floral maturity occurring 30 to 50 years after germination [77,78].

Compared to other studies in which high-throughput sequencing techniques have been used for the development of new microsatellite markers in species with very little information available [78,79], our results are similar. We identified 57 potential SSR markers, of which 11 displayed polymorphism and were used to assess the genetic structure of *B. aethiopum* populations in Benin. We find a low genetic diversity, with an average He value (0.354) that is substantially below those reported for *Borassus flabellifer* [46] and for other non-timber forest products such as *Khaya senegalensis* (He = 0.53; [80] and *Phyllanthus sp* [81]. The positive F value that we observed in the majority (6 out of 9) of populations in the present study indicates an overall deficiency of heterozygotes across population. This deviation from the Hardy-Weinberg equilibrium (HWE) might reflect poor gene flows through pollen and seed dissemination, leading to crosses between related individuals. Accordingly, our data reveal limited genetic distances among populations, with values lower than those reported for others palm species. Indeed for *B. flabellifer*, genetic distance ranged from 0.716 to 0.957 [82] and among natural oil palm accessions an average of 0.769 was observed [84]. Both our Fst values and AMOVA analysis point to intra-population differentiation as being the main source of genetic variation. As illustrated by the agreement between our PCoA and Bayesian analyses, Beninese *B. aethiopum* populations cluster globally according to geographic distances between the collection sites. However, among the nine populations studied, the population from Savè appears to be the most diversified (He= 0.451) and constitutes an exception to this general distribution. This site located in the Sudano-Guinean transition zone of Benin is currently the most active for the production of fan palm hypocotyls, and it acts as a supplier for the whole national territory (VK Salako, personal communication), suggesting that it might be the largest population of *B. aethiopum* in the country. Moreover, our sampling of Savè individuals appear to be genetically distinct from palms belonging to other populations of the Central region and closer from those of the Northern region, despite the important geographical distances involved with the latter case. We postulate that seed dispersion by elephants might have played a major role in the observed pattern of genetic diversity and explain the singularity observed in Savè. As a matter of fact, [31,32] detected the presence of *B. aethiopum* seeds in elephant dungs and hypothesized that elephants may have played important role in the seed dissemination for this species through fruit consumption and long-distance herd migrations. In support to this assumption, Savè is part of a continuous forest corridor connecting with the Northern region that was used by elephants in their migrations. Up until 1982, the seasonal occurrence of the animal has been reported in the Wari-Maro forest of Central Benin [84].

The use of the specific microsatellite markers developed in this study from genomic sequencing of *B. aethiopum* appears to be efficient to assess the genetic diversity and population structure of this species. These microsatellite loci with respect to our results represent potential molecular marker set that can be used to elucidate the genetic diversity of *B. aethiopum* in other African countries. Additionally, and provided that genome divergence is not too extensive to allow marker transferability, our SSR markers may also been used in a palm species that belongs to the same genus and that is reported to share parts of its distribution area, namely *Borassus akeassii* B.O.G., which has long been confused with *B. aethiopum* due to its similar morphology [85]. High-throughput sequencing proves to be a good, fast and effective way to develop new microsatellite markers especially for plant species without published molecular data. The increasing availability and affordability of this technology makes it possible, both technically and financially, to overcome the difficulties arising in case studies such as ours, where marker transfer has proven to be limited or ineffective. To our knowledge, the data presented in the present article constitute the first sizeable molecular resource available for the African fan palm, which we have made available to the scientific community at large in order to facilitate the implementation of an increasing number of studies on this palm species. We have also performed the first analysis of the genetic diversity of *B. aethiopum* in an African country, which we see as a first step towards the elaboration of an evidence-based strategy for sustainable resource management and preservation in Benin. As a complement, the acquisition of agro-morphological data and the characterization of processes regulating the reproductive development of the species are currently under way. Beyond that, we also aim to extend our analysis of *B. aethiopum* diversity to the West African sub-region, and leverage the data acquired to improve knowledge of both other species within the *Borassus* genus, and of palms diversity as a whole.

## Acknowledgements

The Authors wish to thank Emira Cherif, Carina Castillo and Frédérique Aberlenc (IRD, UMR DIADE, Montpellier, France) for sharing their unpublished microsatellite markers data for transferability tests, and Dr Jaume Pellicer (Comparative Plant & Fungal Biology, Jodrell Laboratory, Royal Botanical Gardens, Kew, UK), for allowing us to use the palm flow cytometry data.

## Ethics statement

In accordance with the Nagoya Protocol on Access and Benefit Sharing (ABS), a field permit allowing access and non-commercial use for research purposes of the plant material used in the present study has been submitted to the competent national authority (Direction Générale des Eaux, Forêts et Chasse/Ministère du Cadre de Vie et du Développement Durable, Benin).

## Supporting information captions

**S1 Table: List of sampled *B. aethiopum* individuals.**

M, F: = male or female palm, respectively.

All geographic coordinates are provided as North from the Equator (latitude) and East from the Greenwich meridian (longitude), respectively.

(DOCX)

**S2 Table: List and characteristics of putative microsatellite loci identified in the genome of *B. aethiopum* through QDD analysis**

(XLSX)

